# Disruption of Zika virus xrRNA1-dependent sfRNA1 production results in tissue-specific attenuated viral replication

**DOI:** 10.1101/2020.09.10.281949

**Authors:** Hadrian Sparks, Brendan Monogue, Benjamin Akiyama, Jeffrey S. Kieft, J. David Beckham

## Abstract

Zika virus (ZIKV), like other flaviviruses, produces several species of sub-genomic RNAs (sfRNAs) during infection, corresponding to noncoding RNA fragments of different lengths derived from the viral 3’ untranslated region (UTR). Over the course of infection, these sfRNAs accumulate in the cell as a result of incomplete viral genome degradation of the 3’UTR by host 5’ to 3’ exoribonuclease (Xrn1). The halting of Xrn1 in the 3’UTR is due to two RNA pseudoknot structures in the 3’UTR termed exoribonuclease-resistant RNA1 and 2 (xrRNA1&2). Studies with related flaviviruses have shown that sfRNAs are important for pathogenicity and inhibiting both mosquito and mammalian host defense mechanisms. However, these investigations have not included ZIKV and there is very limited data addressing how sfRNAs impact infection in a whole animal model or specific tissues. In this study, we rescued a sfRNA1-deficient ZIKV (X1) by targeted mutation in the xrRNA1 3’ UTR structure. We found that virus which lacks the production of the largest ZIKV sfRNA species, sfRNA1. Using the X1 virus to infect adult *IFNAR1^-/-^* mice, we found that while the lack of sfRNA1 does not alter ZIKV replication in the spleen, there is a significant reduction of ZIKV genome replication in the brain and placenta compared to WT ZIKV infection. Despite thee attenuated phenotype of the X1 ZIKV, mice develop a robust neutralizing antibody response. We conclude that targeted disruption of xrRNA1 results in tissue-specific attenuation while still supporting robust neutralizing antibody responses. Future studies will need to investigate the tissue-specific mechanisms by which ZIKV sfRNAs influence infection and may utilize targeted xrRNA mutations to develop novel attenuated flavivirus vaccine approaches.

## Introduction

Arthropod-borne flaviviruses such as West Nile virus (WNV), dengue virus (DENV), and Japanese encephalitis virus (JEV) are important human pathogens. More recently, the Zika virus (ZIKV) pandemic and continued increase of tick-borne encephalitis virus infections emphasize the ongoing need to better define the mechanisms of flavivirus pathogenesis. The flavivirus genome is a positive-sense, single-stranded RNA encoding a 5’ untranslated region (UTR), a single open reading frame (ORF), and a highly structured and conserved 3’ UTR. One of these conserved RNA structures in the 3’UTR is known to inhibit degradation of viral RNA by host exoribonucleases [1]. These exoribonuclease-resistant RNAs (xrRNAs) form a pseudoknot structure dependent on a complex tertiary structure which protects the remaining 3’ UTR and results in the accumulation of sub-genomic flaviviral RNAs (sfRNAs) during infection [1–3]. One or more xrRNAs and the biogenesis of sfRNAs has been identified for several flaviviruses including WNV, Murray Valley encephalitis virus (MVE), JEV, DENV, and ZIKV [1–5].

The specific role of sfRNAs during infection is not well-defined, especially *in vivo*. Recent *in vitro* studies have indicated that sfRNAs are important for limitation of cytoplasmic mRNA decay, induction of cell apoptosis, control of the mammalian host type I interferon (IFN) response, host switching, and inhibition of RNAi and Toll receptor pathways in arthropod hosts [2, 4, 6–11]. Due to lack of structural data for flavivirus xrRNAs, many studies have relied on extensive modifications in the 3’ UTR to completely eliminate the biogenesis of all sfRNA species. With the recent publication of detailed structural data for the first of two xrRNAs in ZIKV (xrRNA1), targeted mutations can now be utilized to disrupt RNA tertiary structure independent of major sequence changes in the 3’ UTR [1]. Using this approach, we made specific mutations in xrRNA1 that disrupt the tertiary structure without significantly altering the 3’UTR sequence.

Our approach allows us to evaluate the function of individual sfRNAs while maintaining the sequence of sfRNAs. This strategy limits the potential for off-target effects due to significant sequence or structural alteration of the 3’ UTR. Both, WNV and DENV produce multiple sfRNA species, largely in correlation to the number of xrRNAs encoded in the 3’ UTR. Similarly, ZIKV generates two consistently observed species of sfRNAs (sfRNA1 and sfRNA2) due to the presence of two xrRNA structures (xrRNA1 and xrRNA2) within the 3’ UTR. Despite this, only limited data has been presented investigating the role of individual sfRNA species. [2, 11, 12]. Perhaps more importantly, the role of sfRNA production during ZIKV infection in the mouse model of disease is not known.

Here we use our previous structural data of ZIKV xRNA1 tertiary structure to generate an infectious ZIKV clone in which a discrete, single nucleotide mutation eliminates the production of sfRNA1 without significant changes to the 3’ UTR sequence or structure. We show that loss of sfRNA1 does not impact vial growth in mammalian cells nor limits infection in mosquito cells. In type I interferon knockout mice, (*Ifnar1^-/-^*), ZIKV clones with the xrRNA1 mutation exhibited significantly decreased viral growth in the brain and placenta while still producing a robust neutralizing antibody response. These data show that sfRNA1 production plays a tissue-specific role in support of viral replication.

## Results

### Development of an infectious ZIKV X1 mutant

As previously described, a highly conserved cytosine (nt 10415) located in the P2’ region of ZIKV xrRNA1 secondary structure is necessary for anti-exoribonuclease activity (Figure 1A). [1, 3, 5] In the tertiary structure of xrRNA1, this cytosine forms several bonds that stabilize the phosphate backbone kink essential to inhibiting degradation by host exoribonucleases (Figure 1B). Replacing C10415 with a guanine would be predicted to disrupt the tertiary structure of xrRNA1 without significantly altering the sequence (Figure 1C). We used site-directed mutagenesis to manipulate an infectious cDNA clone of Puerto Rican strain PRVABC59 and produce an infectious clone of the X1 mutant. Sanger sequencing confirmed the presence of the X1 mutation. To rescue infectious virus, Vero cells were transfected with RNA transcribed from the PRVABC59 clone with either the X1 mutation (X1) or no mutations (WT). Following transfection, we determined ZIKV genome copies present in the supernatant of transfected cells to evaluate X1 virus replication. We found that both WT and X1 transfections produced a mean of 2.3X109 viral genome copies per μL, respectively with no significant difference between the WT and X1 clones (Figure 1E). Additionally, there was no significant difference in the amount of infectious WT and X1 virus as measured by a focus forming unit assay (FFU). In this assay, we found that the transfection of ZIKV produced 1.7X106 FFU per mL of supernatant while X1 produced 8.7X105 FFU per mL (Figure 1F). These results indicate that the X1 mutation did not significantly alter viral production after transfection and rescue of infectious clones.

**Figure 1.**
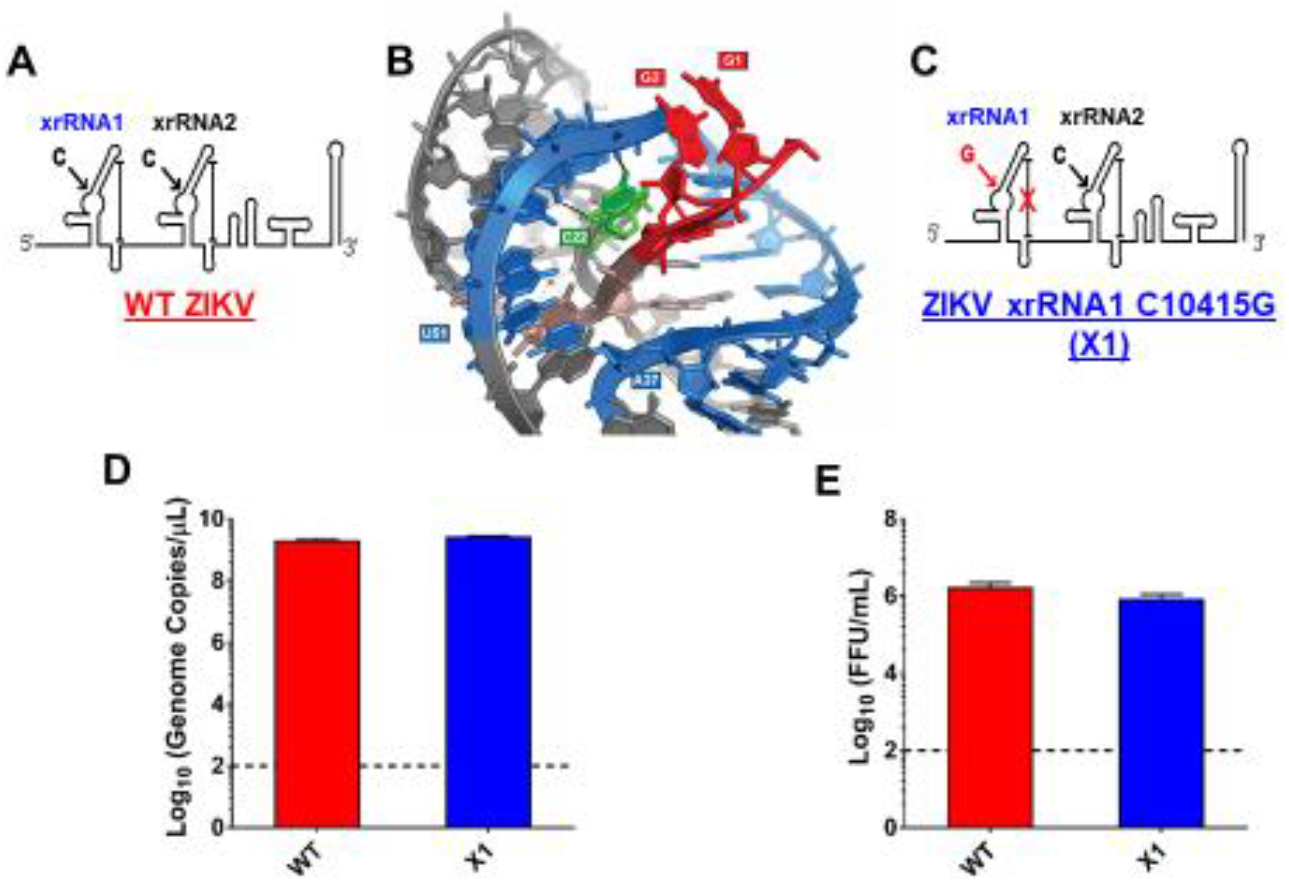
Development of infectious ZIKV with discreet xrRNA1 structural mutation. (A) The 3’ UTR of ZIKV encodes two host exoribonuclease resistant RNA structures (xrRNAs). (B) The tertiary structure of ZIKV xrRNA1 has been previously described, revealing that a single cytosine at position 10415 (green) of the crystalized RNA fragment is necessary for stabilizing the phosphate backbone kink (blue) which protects the viral 3’ UTR (red) from exoribonuclease degradation. We produced a ZIKV mutant (called X1) in which the cytosine at position 10415 of xrRNA1 has been replaced with a guanine (C), weakening the tertiary structure of the RNA. A wild type (WT) ZIKV clone with no mutations (A) was also produced alongside the X1 mutant as a positive control. The viral genomes of either X1 or the WT clone were transfected into Vero cells. At 12 days post transfection, the amounts of both viral genome (D) and infectious virus (E) rescued from the X1 transfection were comparable to those rescued from cells transfected with WT ZIKV RNA (*n*=6).

### X1 mutation does not significantly alter viral growth *in vitro*

To elucidate the sfRNA phenotype of our X1 infectious clone, the production of sfRNAs during infection was evaluated by northern blot of total RNA from infected A549 cells (Figure 2A) [1, 3]. When compared to cells infected with WT ZIKV, we determined that X1 virus only produces sfRNA2 and lacks any observable sfRNA1 production (fig 2A). Interestingly we observe that an sfRNA3 fragment becomes prominent upon loss of xrRNA1. This sfRNA3 band is likely generated by additional unknown mechanisms within the infected cell, independent of the ability for xrRNA1 to resist exonucleolytic decay. Next, we inoculated human A549 cells and U4.4 cells (*Aedes albopictus* cells) with clone-derived X1 and WT viruses (Figure 2B-E). Confluent cells were infected at an MOI of 0.1 and supernatant was harvested at 0, 24, 48, and 72 hours post infection (HPI) to measure viral genome and infectious virus using RT-PCR and FFU, respectively. We found that extracellular ZIKV genome increased with similar kinetics over 72 hours following inoculation with X1 and WT clone-derived virus in A549 cells (2-way ANOVA, *p*=0.9423 Figure 2B). Using viral titer as measured by FFU, we also found no significant difference in infectious virus titers over 72 hours when comparing clone-derived X1 and WT viruses (2-way ANOVA, *p*=0.4603 Figure 2C). However, in U4.4 cells, we observed that both X1 genome copies (2-way ANOVA, *P*=0.9134 Figure 2D) and infectious viral titer (2-way ANOVA, *P*=0.4782, Figure 2E) increased during infection, maintaining similar growth kinetics between time points.

**Figure 2.**
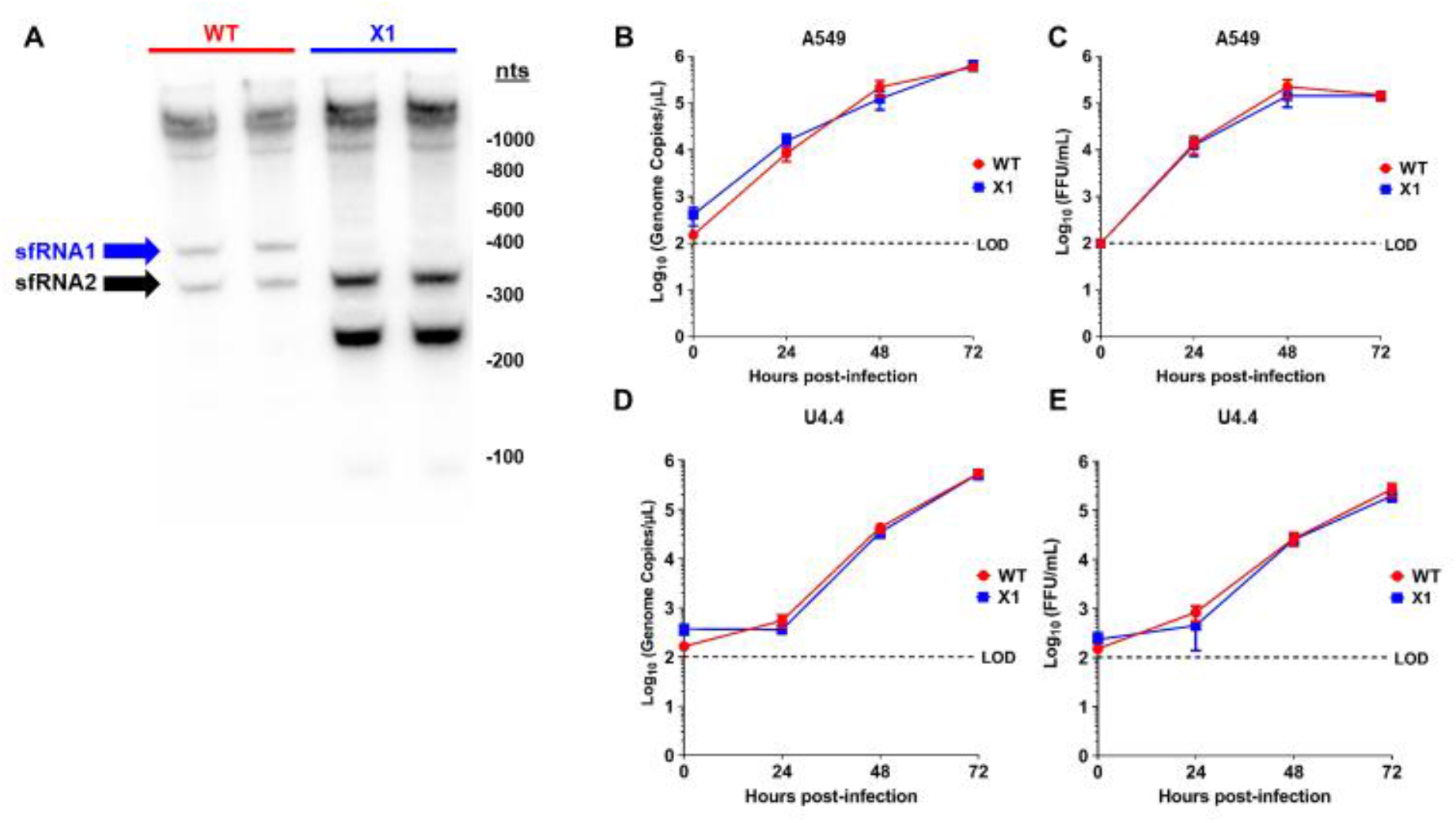
*In vitro* sfRNA production and viral growth kinetics of X1 compared to WT ZIKV Clone. (A) A549 cells were infected with either X1 or WT Clone at an MOI of 1. At 48 hpi, cellular RNA was collected and the presence of ZIKV sfRNAs was detected via Northern blot using a ZIKV 3’UTR-specific probe. To analyze viral growth kinetics, human A549 cells (B-C) or Aedes albopictus U4.4 cells (D-E) were infected with X1 or WT at an MOI of 0.1. At 0, 24, 48, and 72 hpi, supernatant was collected and used to measure either extracellular viral RNA via RT-qPCR (B, D) or infectious virus via FFU assay (C, E). (B-E) Dashed lines represent the limit of detection (LOD). Error bars indicate standard error of the mean for six replicates across two independent experiments. (*n*=6, NS by two-way ANOVA).

These data demonstrate that the clone-derived X1 ZIKV results in loss of sfRNA1 expression with a single nucleotide substitution, and loss of sfRNA1 production does not significantly alter viral growth in cultured A549 cells.

### X1 ZIKV is attenuated in adult *Ifnar1^-/-^* mice and produces neutralizing antibody responses

Since the WT ZIKV infectious clone is attenuated in wild-type mice [13], we compared acute infection with X1, WT ZIKV, or the original clinical isolate PRVABC59 in adult *Ifnar1^-/-^* mice. This model has been established as an important and relevant model of acute ZIKV infection in mice [14]. Moreover, infection of *Ifnar1^-/-^* mice was previously shown to rescue the virulence of a WNV clone in which production of sfRNA1 and sfRNA2 was eliminated through several deletion and substitution mutations [10]. *Ifnar1^-/-^* mice aged 5-7 weeks were inoculated by intraperitoneal (IP) injection with 104 FFU of X1, WT, or PRVABC59 (*n*=5). To determine early viral load, we performed a retro-orbital bleed at 2 days post-infection (DPI) and measured the amount of ZIKV genome present in the serum via RT-qPCR. We found that all mice infected with PRVABC59 or WT ZIKV had quantifiable viral genome (102-103 copies/μL serum) present with no significant difference between the two infections (Figure 3A). However, only 2 of the 5 mice infected with X1 had measurable viral genome in the serum at 2 DPI (Figure 3A). Additionally, *Ifnar1^-/-^* mice inoculated with the X1 mutant virus gained weight during acute infection with an average increase of 3.8% at 6 DPI (Figure 3B). This weight gain was significantly higher than mice infected with WT ZIKV or PRVABC59, which lost 3% and 1% body weight respectively at 6 DPI (2-way ANOVA, *P*<0.05, Figure 3B). While some mice exhibited up to 6% weight loss during these studies, no difference in mortality was observed in any experimental group.

**Figure 3.**
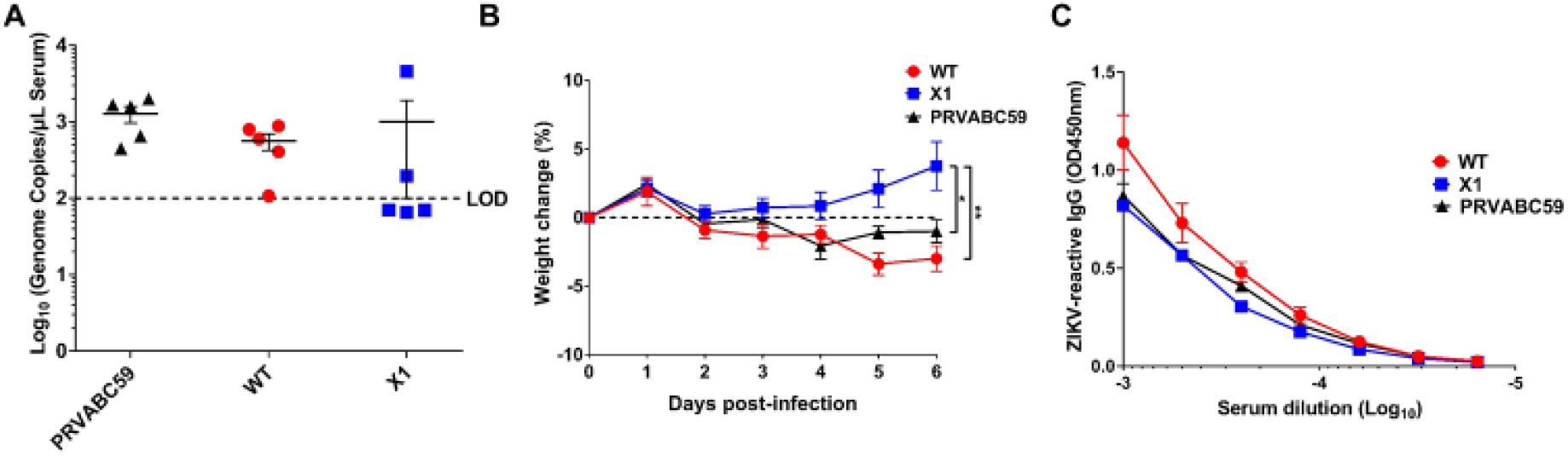
X1 infection compared to WT Clone and ZIKV PRVABC59 in adult *IFNAR1-/-* mice. Male and female *IFNAR1-/-* mice 5-7 weeks in age were infected IP with 1e4 FFU of X1, WT ZIKV Clone, or PRVABC59 virus. (A) Serum samples were collected via RO bleed at 2 dpi to quantify early viral infection. RNA was isolated from the sera and used to detect ZIKV genome via RT-qPCR. Dotted line represents the limit of detection (LOD), no significant differences were found using Mann-Whitney tests to compare between viral infections (*n*=5). (B) The weight of infected mice was monitored during acute infection and shown here as percent weight change relative to baseline set at 0 dpi. Dotted line symbolizes 0% weight change. Two-way ANOVA was used to make multiple comparisons at each time point, asterisks representative of the following: *,X1 vs PRVABC59: *P*=0.0464 at 5 dpi and *P*=0.0014 at 6 dpi. **,X1 vs WT: P<0.0001 at 5 and 6 dpi. (*n*=5). (C) Serum was collected via cardiac stick at 20 dpi to detect a ZIKV-reactive antibody response. ZIKV-reactive IgG was detected by indirect ELISA using ZIKV virions as antigen and donkey α mouse IgG HRP conjugate as detecting antibody. Data shown as optical density (OD) at various serum dilutions, normalized with uninfected mouse sera (*n*=2, NS).

We also assessed the antibody response elicited by these infections at 20 DPI via indirect ELISA using purified ZIKV particles as antigen and a donkey anti-mouse IgG HRP conjugate antibody to detect ZIKV-reactive IgG in infected mouse sera by ELISA. We found that despite the attenuated phenotype of X1 ZIKV, the X1 ZIKV-inoculated adult *Ifnar1^-/-^* mice exhibited equivalent production of ZIKV-reactive IgG antibodies when compared to WT inoculated adult *Ifnar1^-/-^* mice (Figure 3C). Together, these results show that WT ZIKV and PRVABC59 produce similar acute infections while X1 exhibits attenuation while generating an antibody response comparable to WT ZIKV and PRVABC59 in *IFNAR1-/-* mice.

### X1 is attenuated in the CNS tissue of adult *Ifnar1^-/-^* mice

Because X1 exhibited attenuated acute *in vivo* infection despite the lack of a functional type I IFN response, we sought to determine if this attenuation was more clearly observed in different tissues. Specifically, we investigated the impact of sfRNA1 deficiency on ZIKV infection of the brain in the absence of a type I IFN response. We inoculated adult *Ifnar1^-/-^* mice with 104 FFU (IP inoculation) of WT or X1 (*n*=6) then collected brain and spleen from both groups at 6 DPI. We found that X1 exhibits differential patterns of infection dependent on tissue. The spleens of *Ifnar1^-/-^* mice (5-7 weeks old) infected with either WT or X1 exhibited similar ZIKV genome copies at 6 DPI (1.6X104 genome copies/mg tissue, Mann-Whitney Test, *P*=0.3939, Figure 4A). However, in the brain we found that X1 exhibited a 78% decrease in genome copies per mg of tissue compared to WT (Figure 4B). This decrease was significantly different and replicated across multiple repetitions of the experiment (Mann-Whitney Test, *P*=0.0022, Figure 4B). To further define this tissue-specific pattern of X1 ZIKV infection, we expanded the time frame of sample collection to include 4 and 8 DPI. Despite the differences in growth in the brain, we observed that both X1 and WT genome in the spleen peaked early during infection at 4 DPI with a mean of 3.6X105 genome copies per mg of tissue (Figure 4C). Indeed, no significant differences in viral genome in the spleen were seen between X1 ZIKV and WT ZIKV at all time points (Figure 4C). Further reinforcing our findings at 6 DPI, we found a two-fold decrease in X1 ZIKV viral genome in brain tissue compared to WT ZIKV during early (4 DPI) and later stages (8 DPI) of acute infection (*p=0.003, Two-way ANOVA, Figure 4D). Overall, X1 ZIKV is able to replicate the infection kinetics of WT ZIKV in the spleens of *Ifnar1^-/-^* mice but exhibits tissue-specific restriction in viral growth in brain tissue. These data indicate that the attenuation of X1 infection is specific to tissues and that this attenuation is not simply due to a temporal lag in X1 ZIKV infiltration of the CNS.

**Figure 4.**
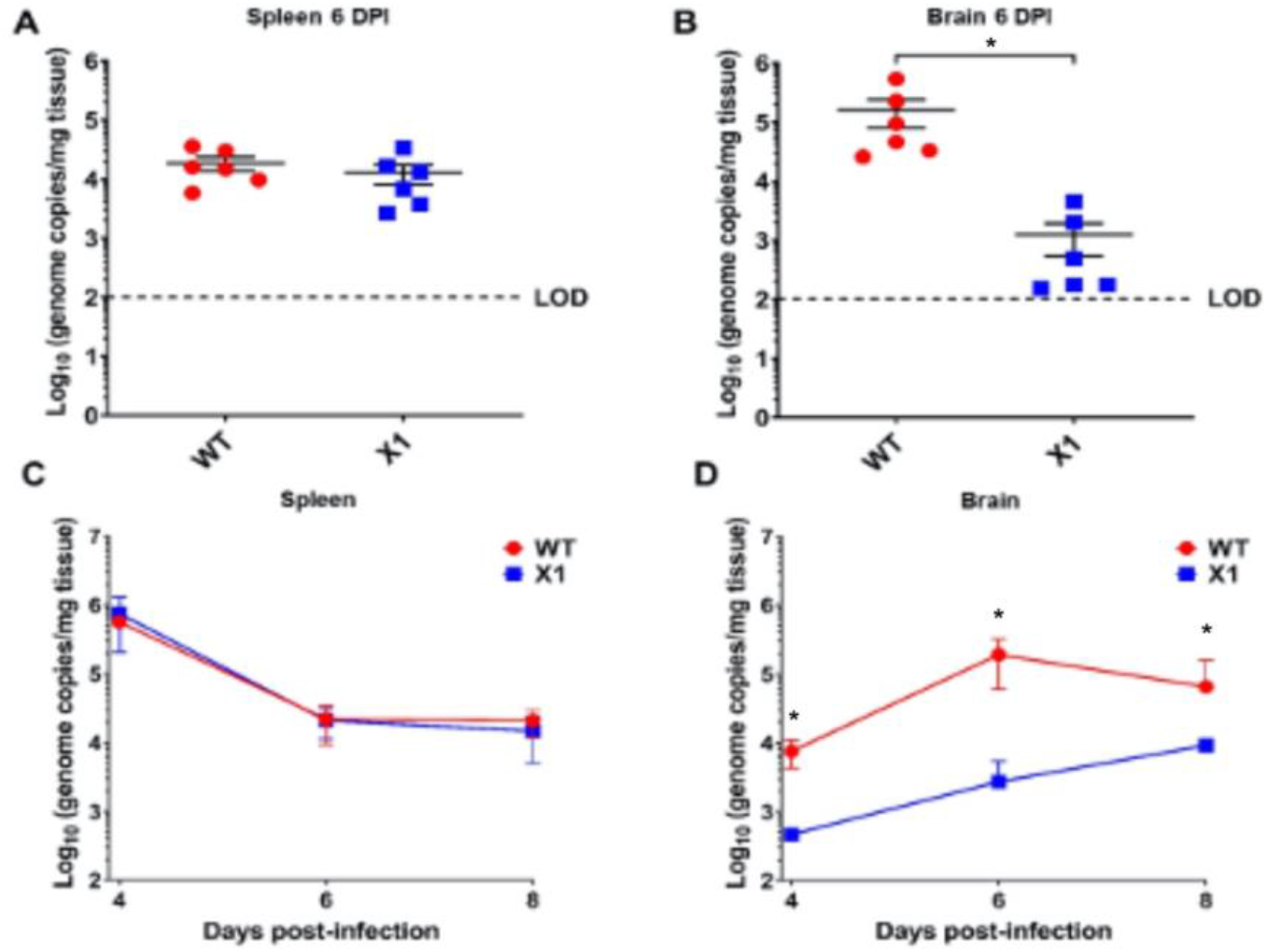
X1 ZIKV exhibits tissue-specific attenuated viral replication in *IFNAR1-/-* mice. Male and female mice were infected with 1e4 FFU (IP) of either WT or X1 virus. At the indicated days post-infection, mice were sacrificed and perfused with PBS before harvest of spleen (A, C) and brain tissue (B, D). Total RNA was isolated from the tissue and used to quantify ZIKV genome via RT-qPCR. Dotted line represents the limit of detection (LOD). Error bars indicate standard error of the mean for six replicates across two independent experiments. [*n*=6 per group, \**P*=0.003 by Mann-Whitney test (B) and two-way ANOVA (D)].

### X1 ZIKV replication is attenuated in the placenta of pregnant *Ifnar1^-/-^* mice

ZIKV vertical transmission to the placenta and fetal tissue are important complications of infection. To further define the impact of the X1 mutation on *in vivo* infection, we characterized X1 ZIKV infection in a pregnant mouse model. Superovulation was used to induce timed pregnancy in *Ifnar1^-/-^* dams mated with *Ifnar1^-/-^* sires (*n*=5, Figure 5A). Dams were infected at embryonic day 6.5 (E6.5) with X1, the clone-derived WT as a positive control, or HBSS to serve as an uninfected control when evaluating fetal outcome via reabsorption. We assessed fetal outcome in these groups at E13.5 (7 DPI) and found that infection with X1 did not induce fetal reabsorption to the extent WT infection had (Figure 5B). ZIKV genome quantified with RT-qPCR of RNA isolated from maternal spleen (Figure 5C) and brain tissue (Figure 5D) showed patterns of X1 and WT infection similar to those seen in adult *Ifnar1^-/-^* mice (Figure 4). No measurable viral genome was detected in the heads of fetal mice from either infection group at this time point (Figure 5E). However, we found that both WT and X1 infected the placenta, and WT-infected placenta exhibited significantly higher ZIKV genome copied, 105 copies/mg tissue, than X1 infected placental tissue, 103 copies/mg tissue (Figure 5F). These findings show for the first time that targeted mutation of xrRNA1 to eliminate sfRNA1 production results in tissue specific attenuation of ZIKV replication in the placenta.

**Figure 5.**
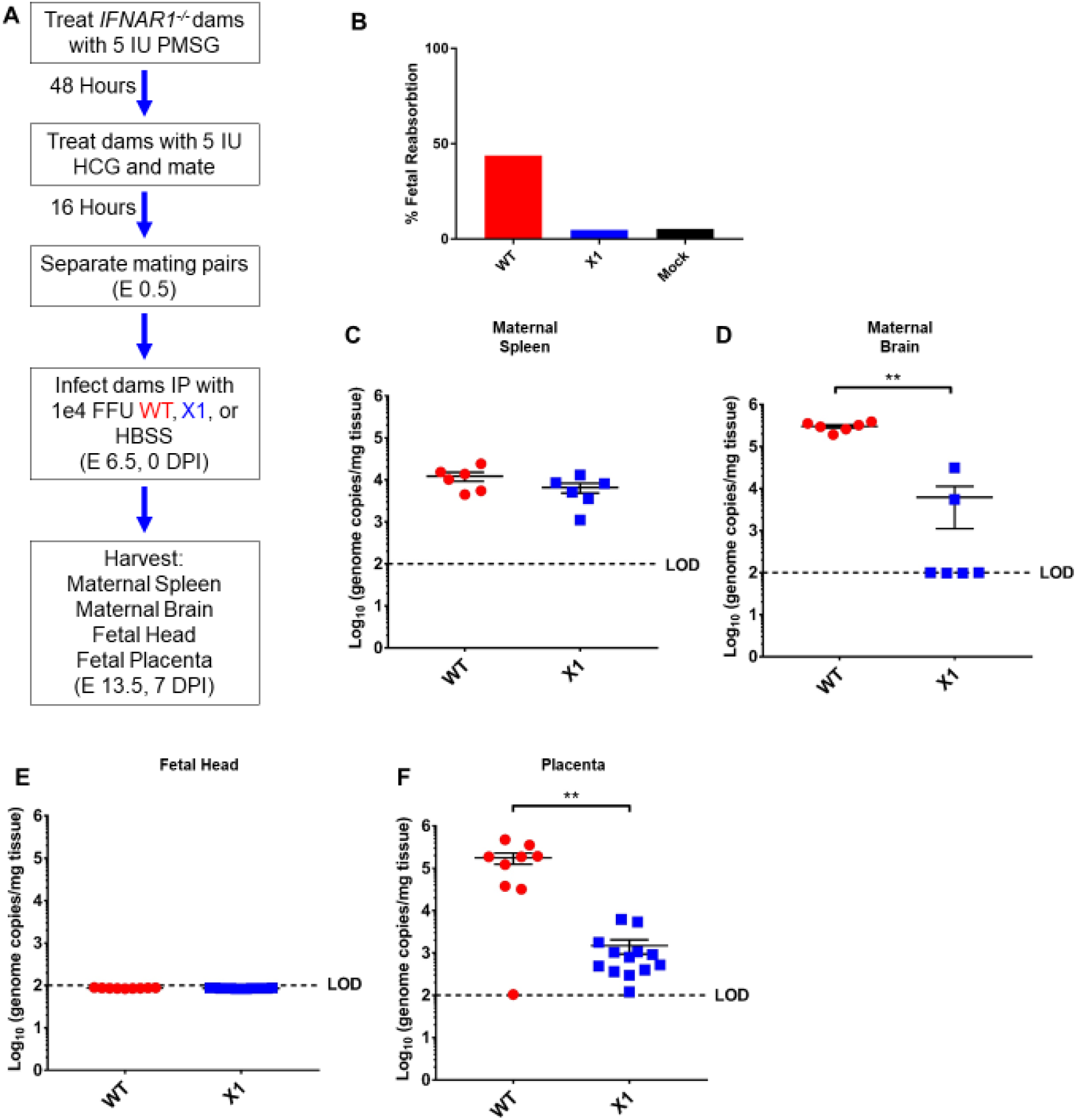
Fetal outcome and viral burden in pregnant *IFNAR1-/-* mice infected with either WT or X1 ZIKV. (A) Superovulation was induced in 10-12 week old *IFNAR1-/-* dams which were then mated with *IFNAR1-/-* sires. Dams were then infected IP with either 1e4 FFU of WT ZIKV clone, the X1 mutant, or 100 ul HBSS at E 6.5. At E 13.5, 7 DPI dams were sacrificed and perfused with PBS. Fetal outcome was assessed and is shown % fetal reabsorption (B). Maternal spleen (C), maternal brain (D) were collected to assess viral burden in these tissues by RT-qPCR of ZIKV genome (*n*=6, \*\**P*<0.01 via Mann-Whitney Test). Fetal head (E), and placenta (F) were also collected to assess viral burden via detection of ZIKV genome (WT *n*=9, X1 *n*=13, \*\**P*<0.01 via Mann-Whitney Test). Fetal data representative of 9-13 fetuses from 3 pregnant dams for each infection. Dotted line shows limit if detection (LOD) for the assay, error bars indicate the standard error of the mean for these experiments.

### Attenuated X1 ZIKV generates a strong neutralizing antibody response in Transgenic STAT2 mice

Use of the *Ifnar1^-/-^* mouse model has provided us with important insights into the pathogenesis of our X1 mutant ZIKV, but this is an immunocompromised model which limits interpretation of attenuation due to the loss of type 1 IFN stimulation. The recently established mouse model with a knocked-in human STAT2 gene (*hSTAT2 KI*) provided an opportunity to assess X1 pathogenesis in an immunocompetent animal model [15]. Adult (5-7 weeks old) *hSTAT2 KI* mice were infected via IP with 10_4_ FFU of either X1, WT ZIKV, or 100 μL of HBSS as a mock infection control (*n*=6). There was no detectable viral genome in the serum of infected animals at 2 DPI and no weight loss was observed during acute infection with either WT or X1 ZIKV (Figure 6A). At 20 DPI, we found that ZIKV-reactive IgG antibodies were detectable in the serum of both WT ZIKV and X1 infected mice (Figure 6B). With an average OD450nm measurement of 0.56, ZIKV X1 infected mice exhibited significantly higher levels of IgG than those infected with WT ZIKV (OD450nm 0.38, 2-way ANOVA, *P*≤0.002, Figure 6B). To determine if serum from infected animals could neutralize WT ZIKV in comparison to mock infected animals, we performed a focus-forming unit reduction neutralization test (FRNT). Serum from mock infected mice did not neutralize or reduce the infectivity of WT ZIKV virions (Figure 6C). The serum from all animals infected with either X1 or WT ZIKV was able to reduce ZIKV infectivity with Log10 EC50 values at −2.98 and −2.88 respectively (Figure 6C). These data illustrate that, despite the attenuated phenotype of the X1 mutant in *IFNAR1-/-* mice, X1 virus produced a strong neutralizing antibody response comparable to WT ZIKV in the *hSTAT2 KI* model of ZIKV infection.

**Figure 6.**
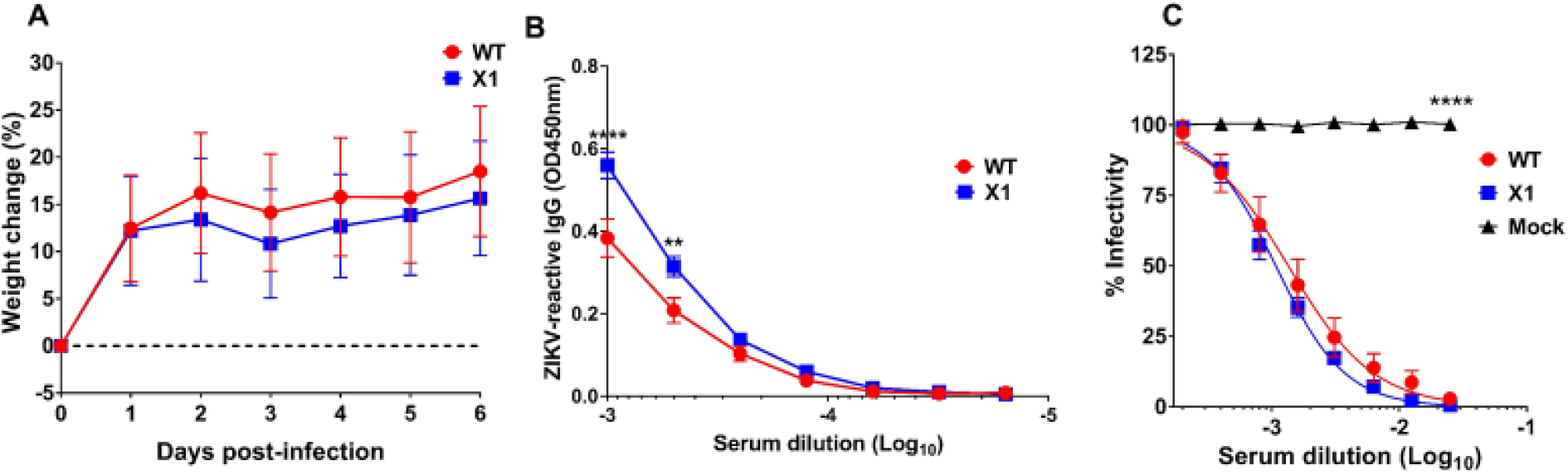
Outcomes of X1 infection compared to WT in HuSTAT2 mice. Male and female HuSTAT2 mice 5-7 weeks in age were infected IP with 1e4 FFU of X1, WT ZIKV Clone, or HBSS as a mock infection. (A) Mouse weight was monitored during acute infection and shown here as percent weight change relative to baseline set at 0 dpi. Dotted line symbolizes 0% weight change with no significant differences detected via two-way ANOVA. (B) Serum was collected via cardiac stick at 20 dpi to detect a ZIKV-reactive antibody response. ZIKV-reactive IgG was detected by indirect ELISA using ZIKV virions as antigen and donkey α mouse IgG HRP conjugate as detecting antibody. Data shown as optical density (OD) at various serum dilutions, normalized with mock infected mouse sera. (C) Serum from 20 dpi was also used to detect neutralization of ZIKV via focus reduction neutralization test (FRNT). Results are shown here as change in percent infectivity of ZIKV compared to untreated ZIKV. All data representative of six replicates across two independent experiments, error bars indicate standard error of the mean. (*n*=6, **** *P*<0.0001, \*\**P*=0.002 via two-way ANOVA).

## Discussion

We utilized the ZIKV xrRNA1 tertiary structure to construct a mutant with a single nucleotide change that disrupts xrRNA1 structure and eliminates sfRNA1 production without significantly altering the sequence of the ZIKV 3’UTR. We show for the first time that targeted mutation of xrRNA1 results in loss of sfRNA1 production and tissue-specific attenuation of ZIKV replication in brain tissue and placenta tissue of type 1 interferon receptor knockout mice. Despite the attenuated phenotype of X1 ZIKV, type 1 interferon receptor knockout mice and transgenic STAT2 KI mice develop robust antibody responses.

Our work has several strengths that add to our current understanding of the role of xrRNA1-dependent production of sfRNA. Previous studies have shown that sfRNA biogenesis is necessary to limit RNAi and Toll pathway activation in arthropods, to limit mRNA decay in host cell cytoplasm, and to impede antiviral type I IFN responses in mammalian cells [4, 7, 10, 16–19]. These previous studies utilized extensive sequence and structural mutation of the 3’ UTR or total elimination of sfRNA1 and sfRNA2 production. Our approach utilized information of the xrRNA1 structure to make mutations that disrupt tertiary structure without significantly altering the sequence of the 3’UTR. Thus, our findings are directly related to the structure-function relationships of xrRNA1 and sfRNA1 production.

Additionally, the scope of the current published literature includes little information on the role of xrRNA1-dependent sfRNA1 production in the pathogenesis of flavivirus infections in animal models. Early investigations with sfRNA1-deficient WNV found that, while this mutation did attenuate overall infection of immunocompetent mice, there was no alteration of viral load in the brain or other tissues [2]. Here, we show for the first time that ZIKV xrRNA1 mutation results in tissue specific attenuation in the brain and placenta in type I IFN receptor knockout mice. Previous studies have shown that type II and type III IFN responses are critical to the control of acute RNA virus infections in the brain and placenta, respectively [20–23]. Recent studies investigating the antiviral response to alphavirus infection of the central nervous system (CNS) have identified IFN-γ and the type II IFN response as being vital for the control of RNA viruses [20, 24]. Given, that we found differences in type 1 IFN deficient mice, future studies evaluating the interactions between xrRNA1 mutant ZIKV and type II and III IFN responses will be critical to fully understand the mechanisms of the sfRNA1 function in different tissues.

We also found a novel infection phenotype in pregnant *Ifnar1^-/-^* mice. Numerous mouse pregnancy infection models have been established to investigate the pathogenesis of congenital Zika virus syndrome as extensively reviewed by Caine et al [25]. However, there have been no findings to date studying sfRNA biogenesis and the impact on pathogenesis in murine pregnancy models of ZIKV. Our initial studies have identified that the sfRNA1-deficient X1 mutant does not efficiently infect the placenta and exhibited decreased fetal reabsorption compared to WT ZIKV infection. Since we did not detect virus in fetal brain tissue, these findings also imply that placental infection plays an important role in ZIKV-induced fetal injury.

The lack of detectable clone-derived WT ZIKV or X1 in the fetal head of infected fetuses identifies an important limitation of this model and corroborates findings that infection with mouse-adapted ZIKV strains produces more consistent fetal infection than common lab strains [26]. Future studies using mouse-adapted ZIKV or insertion of envelope protein virulence determinants as recently described along with our identified xrRNA1 mutations would likely provide more information on the pathogenesis of ZIKV with and without xrRNA1 mutations in fetal tissues [26].

The 3’UTR is a common target for attenuation approaches and vaccine development strategies for flaviviruses though the mechanism for this attenuation is often not well defined [27–31]. Here we have found that ZIKV containing a targeted mutation in the xrRNA1 structure induces a strong ZIKV-specific antibody response in both immune-deficient and immunocompetent mouse models despite the attenuated phenotype. This is especially intriguing given that the xrRNA1 structure altered in our studies is also present in other flaviviruses of global health importance like DENV and WNV [1, 3, 5, 32]. Future studies targeting specific mutations to the xrRNA1 structure in other flaviviruses may provide a novel attenuation approaches for vaccine development in ZIKV as well as other flaviviruses.

Previous studies have shown that sfRNA1 deficiency does not alter growth of WNV or DENV in cultured host cells but does result in reduced CPE and plaque size [6, 12, 33]. Differential expression of sfRNA isotypes assists in host adaptation for some flaviviruses [32, 34, 35]. However, ZIKV consistently produces equal amounts of sfRNA1 and 2 in both mosquito and mammalian host cells suggesting that production of a specific sfRNA species is not crucial for survival in different hosts [1, 35]. We have now shown that sfRNA1-deficient ZIKV X1 virus displayed reduced ability to replicate in the brain and placenta. These data will provide an important starting point for future studies that investigate the tissue-specific role of sfRNA1 and sfRNA2 production during flavivirus infection.

In conclusion, we have developed and characterized an sfRNA1-deficient ZIKV virus through minimal sequence manipulation of a 3’UTR RNA structure and this approach can be utilized to expand understanding of sfRNA function in ZIKV and related flaviviruses. Validation of this model for disrupting sfRNA production can also be applied to ZIKV xrRNA2 or other xrRNA structures for other vector-borne flaviviruses. Further studies examining the role of sfRNA1 and sfRNA2 expression in a tissue specific manner will provide novel insight into the pathogenesis of flavivirus infections and may provide novel attenuation approaches for vaccine development.

## Materials and Methods

### Cell lines and viruses

African green monkey kidney epithelial cells (Vero E6), human lung epithelial cells (A549), and *Aedes albopictus* cells (C6/36) were sourced from the American Type Culture Collection (ATCC). Vero and C6/36 cells were cultured in Eagle’s minimum essential medium (MEM), A549 cells were cultured in Ham’s F-12K medium. Both MEM and Ham’s F-12K medium was supplemented with 1mM sodium pyruvate, 1X non-essential amino acids (100X; ThermoFisher scientific), 100 U/ml streptomycin, 100 μg/ml streptomycin, 10 mM HEPES, and 10% fetal bovine serum (FBS). Dcr-2-expressing *Aedes albopictus* cells (U4.4) were generously provided by Dr. Aaron Brault (CDC, Division of Vector-Borne Diseases) and cultured in Mitsuhashi and Maramorosch insect medium (M&M) supplemented with 10% FBS, 1X non-essential amino acids, 100 U/ml streptomycin, and 100 μg/ml streptomycin. Mammalian-derived cells were maintained at 37 °C with 5% CO_2_ while Aedes-derived cells were maintained at 28 °C with 5% CO_2_. Viruses used in this study include ZIKV Puerto Rico isolate PRVABC59 (science paper), a WT clone derived from PRVABC59, and the X1 mutant ZIKV [13].

### Plasmids and generation of the X1 mutant

Previously described pACYC177 vector plasmids containing ZIKV PRVABC59 genome from either the 5’ UTR to nt 3498 (p1-ZIKV) or from nt 3109 to the end of the 3’ UTR (p2-ZIKV) were used to generate WT ZIKV or the X1 mutant. Primers ZIKV X1 C35G F (5’-TCCCCAAGCTGTGCCTGACTAGCAGGC-3’) and ZIKV X1 C35G R (5’-GCCTGCTAGTCAGGCACAGCTTGGGGA-3’) were used with a QuikChange II XL Site-Directed Mutagenesis Kit (Agilent) to introduce the X1 C10415G mutation into the p2-ZIKV plasmid. The resulting X1 mutant p2-ZIKV, untreated WT p2-ZIKV, and p1-ZIKV plasmids were rescued and amplified via RCA as described [13]. Prior to *in vitro* transcription of the viral genomes, the presence of the C10415G mutation in the X1 mutant was confirmed via Sanger sequencing (Eton Bioscience: San Diego, CA).

### Rescue and propagation of ZIKV

Viral RNA genome for both WT ZIKV and the X1 mutant was in vitro transcribed from ligated plasmid DNA using a HiScribe T7 ARCA mRNA kit (NEB). Approximately 40 μg of the resulting mRNAs were transfected into Vero cells using MessengerMAX lipofectamine transfection reagent (Invitrogen). Supernatant was collected from transfected cells once 50-60% cell clearance was observed and spun down to eliminate cell debris. Clarified supernatant containing the rescued WT or X1 virus was aliquoted stored at −80 °C for later use. Extracellular viral RNA and infectious virus was quantified as detailed below to evaluate the success of viral rescue. Both rescued WT and X1 viruses were subsequently passaged once in C6/36 cells to increase viral titer.

### Virus quantification

The amount of cell-free infectious virus in the supernatant of infected cells was quantified using a focus forming unit assay as previously described [36]. Mouse anti-Flavivirus group antigen antibody, clone D1-4G2-4-15 (Millipore) was used as primary antibody and donkey anti-mouse IgG antibody conjugated with horseradish peroxidase (Jackson Research Laboratories) served as the secondary antibody. Extracellular viral RNA was extracted from the supernatant of infected cells or serum from infected animals using an E.Z.N.A. viral RNA kit (Omega Bio-Tek) and used to quantify viral genome present via the following RT-qPCR protocol. Extracted RNA was reverse transcribed using iScript cDNA synthesis (Bio-Rad). Primers ZikalÜ87 (5’-CCGCTGCCCAACACAAG-3’), Zika1163c (5’-CCACTAACGTTCTTTTGCAGACAT-3’), and FAM-tagged probe Zika1108FAM (5’-AGCCTACCTTGACAAGCAGTCAGACACTCAA-3’) were used in combination with a standard curve spanning 107 copies/reaction to 1 copy/reaction to quantify ZIKV genome copies via qPCR. Transformed data were presented as (log_10_) viral genome copies per microliter.

### Northern blot

Total RNA was isolated from A549 cells infected with either WT or X1 ZIKV and used to evaluate the presence of sfRNAs via Northern blot as previously described [1].

### In vitro viral growth kinetics comparison

2×10^4^ A549 cells or 4×10^4^ U4.4 cells per well were seeded in 24-well plates. Cells were infected with either WT or X1 virus at an MOI of 0.1 for 1 hour then washed with 1XPBS to remove extracellular virus before adding back typical cell growth media. At 0, 24, 48, and 72 hours post infection (HPI) supernatant from infected cells was collected and used to quantify the amount of infectious virus (via FFU assay) or viral genome (via RT-qPCR) present.

### Animal Studies

All animal and infectious disease studies were reviewed by the University of Colorado Institutional Animal Care and Use Committee (IACUC) and Institutional Biosafety Committee. C57BL/6 *Ifnar1^-/-^* mice purchased from Jackson Laboratories and C57BL/6 mice expressing human STAT2 protein (*hSTAT2 KI*) graciously provided by Dr. Michael Diamond (Washington University School of Medicine) were maintained and bred in specific-pathogen-free facilities at the University of Colorado Anschutz Medical Campus animal facility. Animals to be infected were housed in an animal BSL-3 (ABSL-3) laboratory. After infection, mice were observed daily for signs of disease, weight loss, or other terminal indicators until the experiment endpoint.

### Characterization of infection in adult *Ifnar1^-/-^* mice

Similar numbers of male and female 5-7 week old *Ifnar1-/-* mice were intraperitoneally infected with 1×10^4^ FFU of either X1, WT-clone, PRVABC59 ZIKV or 100μL of HBSS as a mock infection. Mouse weight was measured daily until endpoint and serum was collected from a retro-orbital bleed 2 days post infection (DPI) to quantify viremia during early infection via RT-qPCR. At 4, 6, and 8 DPI, a cohort of mice were euthanized with isoflurane and perfused with 20 ml PBS before brain and spleen tissues were collected. Tissues were stored in RNAlater solution (ThermoFisher) at 4 °C until processing. To evaluate the presence of ZIKV-reactive IgG, mice were euthanized with isoflurane at 20 DPI and blood was collected via cardiac puncture. Serum extracted from these blood samples was stored at −80 °C until use.

### Infection in pregnant *Ifnar1-/-* mice

Superovulation was induced in female *Ifnar1-/-* mice aged 10-12 weeks as previously described [37]. Dams were mated with *Ifnar1-/-* sires for approximately 16 hours and then separated (E0.5). At E6.5, dams were infected intraperitoneally with 1×10^4^ FFU of either X1 or WT ZIKV or 100 μL HBSS as a mock infection. 7 DPI on E13.5, mice were sacrificed using isoflurane, perfused with 20 ml PBS, and maternal brain and spleen were collected. Fetal outcome was assessed by number of fetuses and resorptions present and fetal head and placenta were collected to determine viral burden. All samples were stored in RNAlater solution (ThermoFisher) at 4 °C until processing.

### Tissue viral load by RT-qPCR

Tissues collected from infected mice were normalized by weight and mechanically homogenized in TRIzol (ThermoFisher) using a BeadBug instrument (Benchmark Scientific). A standard TRIzol/chloroform RNA isolation protocol was used to reduce lipid contamination [38]. Total RNA was then isolated from the resulting aqueous layer using an E.Z.N.A. total RNA I kit (Omega Bio-Tek). An iScript gDNA clear cDNA synthesis kit (Bio-Rad) was used to reverse transcribe 1000 ng of RNA isolated from infected tissues. The resulting cDNA was used to quantify the amount of ZIKV genome present by qPCR.

### Infection of adult HuSTAT2 mice

Male and female 5-7 week old HuSTAT2 mice were infected intraperitoneally with 1×10^4^ FFU of either X1 or WT ZIKV or 100 μL HBSS as a mock infection. Weight of infected animals was monitored daily. Mice were euthanized with isoflurane at 20 DPI and blood was collected via cardiac puncture. Serum isolated from blood samples was used to determine the amount of ZIKV-reactive IgG present via indirect ELISA as well as viral neutralization by focus reduction neutralization test (FRNT).

### ZIKV-reactive IgG ELISA

Immulon 4HBX 96-well ELISA plates (ThermoFisher) were coated overnight at 4 °C with 200ng/well of ZIKV virus particles diluted in PBS (pH 7.4). Plates were blocked with SuperBlock (ThermoFisher) at room temperature for 1.5 hours then washed once with PBS-T (0.05% Tween 20). Sera from infected animals was serially diluted in PBS-T and incubated on the plate for 1.5 hours at room temperature then washed 3 times with PBS-T. Horseradish peroxidase-conjugated donkey anti-mouse IgG antibody (Jackson Research Laboratories) was diluted 1:4,000 and incubated on the plate for 45 minutes the washed 6 times with PBS-T. TMB substrate (ThermoFisher) was added to the plate and allowed to develop for 3 minutes before 0.3N H_2_SO_4_ was added to stop the reaction. ELISA absorbance was read at 450nm for 0.1 second on a Victor X5 plate reader (PerkinElmer).

### FRNT measurement of serum neutralization

1×10^4^ Vero cells per well were seeded onto opaque Nunc-immuno 96-well plates (MilliporeSigma). Heat-inactivated sera from infected animals was serially diluted and then incubated with WT ZIKV at a concentration of 10 FFU per well at 37 °C for one hour. The resulting serum-treated virus was then used to infect the plated Vero cells for one hour at 37 °C. Cells were then overlaid with a 1:1 mixture of 2.5% Avicel (MilliporeSigma) and complete 2×MEM medium and incubated at 37 °C for 48 hours. After incubation, the cells were fixed with 4% PFA then washed with PBS and stored at 4 °C until staining. The presence of ZIKV in infected cells was detected with a mouse anti-flavivirus 4G2 primary antibody (Millipore) and a donkey anti-mouse IgG horseradish peroxidase-conjugated secondary antibody (Jackson Research Laboratories) as previously described [36]. A nonlinear regression analysis was used to determine the plasma dilution factor at which 50% ZIKV neutralization occurs compared to an untreated control.

### Statistical analysis

GraphPad Prism 7 software (GraphPad Software) was used to analyze all data. Differences were considered statistically significant if *P*<0.05. Specific statistical analysis methods are detailed in the figure legends of experimental results.

## Acknowledgements

We thank Dana Fader and Aaron Massey for research assistance and support during the preparation of this data. We also thank Michael S. Diamond and the Washington University School of Medicine in St. Louis for providing the *hSTAT2 KI* mice.

This work was supported by DOD PRMRP funding (contract W81XWH-17-1-0183) and VA Merit Funding (I01BX003863) and to J.D.B.

We declare that we have no conflicts of interest.

